# DNA methylation landscape of ocular tissue relative to matched to peripheral blood

**DOI:** 10.1101/075077

**Authors:** Alex W Hewitt, Vania Januar, Alexandra Sexton-Oates, Jihoon E Joo, Maria Franchina, Jie Jin Wang, Helena Liang, Jamie E Craig, Richard Saffery

**Author notes:** These authors contributed equally to this work (Alex W Hewitt, Vania Januar). Address for correspondence: Prof Richard Saffery, Murdoch Childrens Research Institute, 50 Flemington Rd, Parkville, Victoria, Australi, 3052, Assoc Prof Alex Hewitt, Centre for Eye Research Australia, 32 Gisborne Street, East Melbourne, Victoria, Australia 3002.

## Abstract

**BACKGROUND:** Epigenetic variation is implicated in a range of non-communicable diseases, including those of the eye. However, investigating the role of epigenetic variation in ocular disease remains problematic as the degree of correlation in epigenetic profile between central (such as the brain or eye) and peripheral tissues (blood or saliva) within an individual remains largely unclear.

**METHODS:** Matched whole blood from the subclavian vein, and whole eyes (N=8) were obtained post-mortem. DNA was isolated from blood, neurosensory retina, retinal pigment epithelium (RPE)/choroid and optic nerve tissue. DNA methylation profiling was performed using the Illumina Infinium HumanMethylation450 platform. Following standard quality control measures a total of 433,768 methylation values common to all samples were available for use in subsequent analysis.

**RESULTS:** Unsupervised hierarchical clustering and principal components analysis revealed tissue of origin as the main driver of variation within the dataset. Despite this, there was a strong correlation of methylation profiles between tissues within each individual. Over 255,000 CpG sites were found to have similar methylation levels (beta <0.2 or beta >0.8) across different tissues in the same individuals, with a further ~16,000 sites having similar methylation profiles across ocular tissues only. Only a small proportion of probes showing interindividual variation in blood, co-varied across blood and eye tissues within individuals.

**CONCLUSIONS:** An improved understanding of the epigenetic landscape of the eye will have important ramifications for regenerative medicine and ongoing dissection of gene-environment interactions in eye disease. Despite a generally high correlation in methylation values irrespective of sample origin, tissue type is the major driver of methylation variation, with only limited covariation between blood and any specific ocular tissue. Caution is warranted when aiming to infer ocular tissue methylation status from blood samples.

## Introduction

The eye is the most specialised sensory organ in the human body. Light enters through the cornea and after passing through the pupil is finely focussed by the crystalline lens onto the neurosensory retina. Initiation of the phototransduction cascade converts the photonic energy into a neural signal, and following a high degree of pre-retinal processing, this signal is transferred via retinal ganglion cells to the brain [1]. Retinal ganglion cells exit the eye thorough the optic nerve to synapse in the mid-brain [1]. Stray light is absorbed by the retinal pigmented epithelium (RPE), which also serves a fundamental role in vitamin A cycling [2]. The high metabolic demand of phototransduction is ameliorated by the cavernous choroidal tissue which is located posterior to the RPE and receives the highest blood flow per tissue volume in the body [2].

Dysfunction of almost any cellular component of the eye can lead to significant visual morbidity. Many ophthalmic diseases are known to have both heritable and environmental pathoaetiological factors, [3] and a greater understanding of the molecular mechanisms of ophthalmic disease have revolutionised therapy [4, 5]. Although much insight into the genetic causes of ocular disease have been gained through well powered genome-wide association and linkage studies, [6] the precise means by which genetic variants and environmental stressors interact remain poorly understood. These dynamic interactions, which have cellular consequences, can be broadly categorised as epigenetic variation. Understanding the epigenetic factors involved in ophthalmic disease may facilitate the development of novel disease screening and therapeutic avenues.

Epigenetic variation has emerged as a major mediator of gene:environment interactions thought to underpin much of human disease. However, unlike genetic variation, epigenetic processes are dynamic both temporally and spatially, with each specific tissue type displaying a unique epigenetic profile. DNA methylation is by far the most widely studied epigenetic process due to its ease of measurement and high degree of stability in biological specimens, and many studies have begun to explore the link between DNA methylation variation and a range of exposures and disease outcomes. Although the field of epigenetic-epidemiology remains in its infancy, some compelling findings have emerged, particularly in relation to the effects of tobacco smoking exposure on DNA methylation profile in blood and buccal cells. A clearer understanding of the methylation landscape of tissues relevant to specific conditions is required in order to gain insights into the relevance of measuring epigenetic profile in a proxy tissue such as blood or saliva [7]. This has already been extensively examined in matched blood and brain [8–10], but has yet to be investigated in matched blood and eye tissue.

The principal aim of this study was to investigate the methylation profiles of specific ocular tissues, and compare this profile to matched peripheral blood in order to assess the utility of blood DNA methylation profile as a proxy for the direct study of eye tissue – not possible in living humans. Herein, we determine the relative importance of ocular tissue methylation specificity at the genome-wide level, and quantify the number of sites that co-vary within an individual across neurosensory retinal, RPE/choroidal and optic nerve tissue.

## METHODS

### Sample collection and processing

Whole blood from the subclavian vein and whole eyes were obtained post-mortem. The donors had no known ophthalmic disease. Donors previously diagnosed with disseminated cancer were excluded. Specimens from eight people were available. These donors were all male and the mean (SD) age was 60.6 (11.3) years (**Supplementary Table 1**).

Blood samples were collected in tubes containing EDTA. All ocular tissue was collected and stored within twelve hours post-mortem. A circumferential pars plana incision was made and the vitreous body discarded. The neurosensory retina was removed and the RPE and choroid were dissected free from the scleral wall. Dura matter was stripped from the anterior segment of optic nerve before storage. Dissected ocular tissue was stored in QIAGEN Allprotect Tissue Reagent (QIAGEN, Hiden, Germany) and DNA extraction was subsequently performed using the QIAGEN DNeasy Blood & Tissue Kit (QIAGEN). Following bisulfite conversion using the Methyl Easy bisulphite modification kit (Human Genetic Signatures, Sydney, NSW, Australia), samples were hybridized to Illumina Infinium HumanMethylation450 (Illumina Inc, San Diego, CA, USA) BeadChips (HM450K) according to the manufacturer’s protocols. Array processing and Beadstudio analysis was conducted through the Australian Genome Research Facility (Melbourne, VIC, Australia).

This study was approved by the human research ethics committee of the University of Western Australia (RA/4/1/4805). Informed consent was obtained from next-of kin or powers of attorney. This study was conducted in accordance with the principles of the Declaration of Helsinki and its subsequent revisions.

### Methylation data preprocessing

Quality control and background correction of the Beadstudio data was performed using the R programming software (version 3.2.1, http://cran.r-project.org/) *ChAMP* Bioconductor package, which utilises the *minfi* package [11]. Normalisation was performed on all samples using the beta– mixture quantile normalisation method, a comparably robust normalisation method which adjusts the beta-value distribution of type II probes to resemble that of type I probes [12]. Probes that had a detection p-value greater than 0.01, probes that were cross-reactive, and probes that potentially contained SNPs of high minor allele frequency [13], were removed from the dataset. As samples were all male, probes that hybridise to sex chromosomes were retained, leaving a total of 433,768 probes. Batch adjustment for samples was performed using the *sva* R package.

### Data processing and statistical analysis

All statistical analysis was performed in R. Unsupervised hierarchical clustering was first performed to investigate the relationships between samples.

Methylation levels between individuals and tissues for all CpG sites were then compared using the Spearman correlation coefficient. CpG sites were categorized as either hypomethylated (β ≤ 0.2), having an intermediate level of methylation (0.2 < β < 0.8), or hypermethylated (β ≥ 0.8), and the overlap of these categories between tissues were investigated.

#### Correlation between tissues

Subsequent analysis utilized probes that were ‘blood variable’, defined as having >5% methylation range within the inner 80^th^ percentile of probes. The association of methylation levels between tissues within individuals was calculated for each CpG site using Spearman correlations. To establish the existence of an association in methylation levels between tissues, the correlation distribution was compared to a null correlation distribution, which was simulated by permuting samples and calculating correlations between unmatched pairs. Paired t-tests with matched samples were used to identify the most similarly methylated probes between tissues.

#### Principal components analysis

To further investigate the structure of variation within the dataset, principal components analysis was performed on normalised M-values. Principal components were analysed for correlation with sample traits (e.g. tissue type, individual), using both linear regression and ANOVA. Probes that are associated with trait-related principal components contribute to the pattern of variation and likely vary across the trait.

As we were mainly interested in inter-individual differences across tissue type, probes that were highly associated with individual-associated principal components (|r| > 0.5) were selected for further analysis. Four principal components were found to vary between individuals only at a p < 0.10 level, accounting for 10.8% of the total variation. Hierarchical clustering was performed with probes related to these principal components, both combined and separately, and with and without the removal of tissue-specific probes. As there was little gene overlap between individual-specific and tissue-specific probes, the removal of tissue-specific probes had no significant effect on clustering. Only one principal component was significantly associated with inter-individual difference at p < 0.05. Probes that correlate with this principal component (PC13) were then compared with probes that co-vary between tissues.

#### Gene annotation and genomic enrichment

Probes were categorised into genomic features and CpG island features according to the annotation file provided for Illumina. Enrichment within each category was investigated for all probes, blood variable probes, blood variable probes that correlate with each eye tissue (|r| > 0.5, p < 0.05), and probes that correlate with individual-related principal components.

#### Gene ontology and Pathway analysis

Ingenutity Pathway Analysis (IPA) was used to investigate cellular pathways subject to DNA methylation variation. Gene ontology enrichment analysis was assessed using the PANTHER Overrepresentation Test (release 20160715) as part of the Gene Ontology Consortium (www.geneontology.org). HM450K probes located in a gene, or within 2kb of the transcription start site showing differential methylation were assigned to the annotated gene. Genes on the HM450K array were used as the background list against which a target list were analysed.

## RESULTS

### Sources of variation within the dataset

Following quality control and data normalization, DNA methylation values from a total of 433,768 probes were available for use in analysis. As anticipated, unsupervised hierarchical clustering revealed tissue type as the main driver of variation within the dataset, blood had the most distinct methylation profile, while RPE/choroid and optic nerve were the most similar (Figure 1). Multidimensional Scaling plot visualisation of the 1,000 most variable probes highlighted the similarity of RPE/choroid and optic nerve methylation profiles relative to retina, with blood the most dissimilar in methylation profile (**Supplementary Figure 1**).

**Figure 1.**
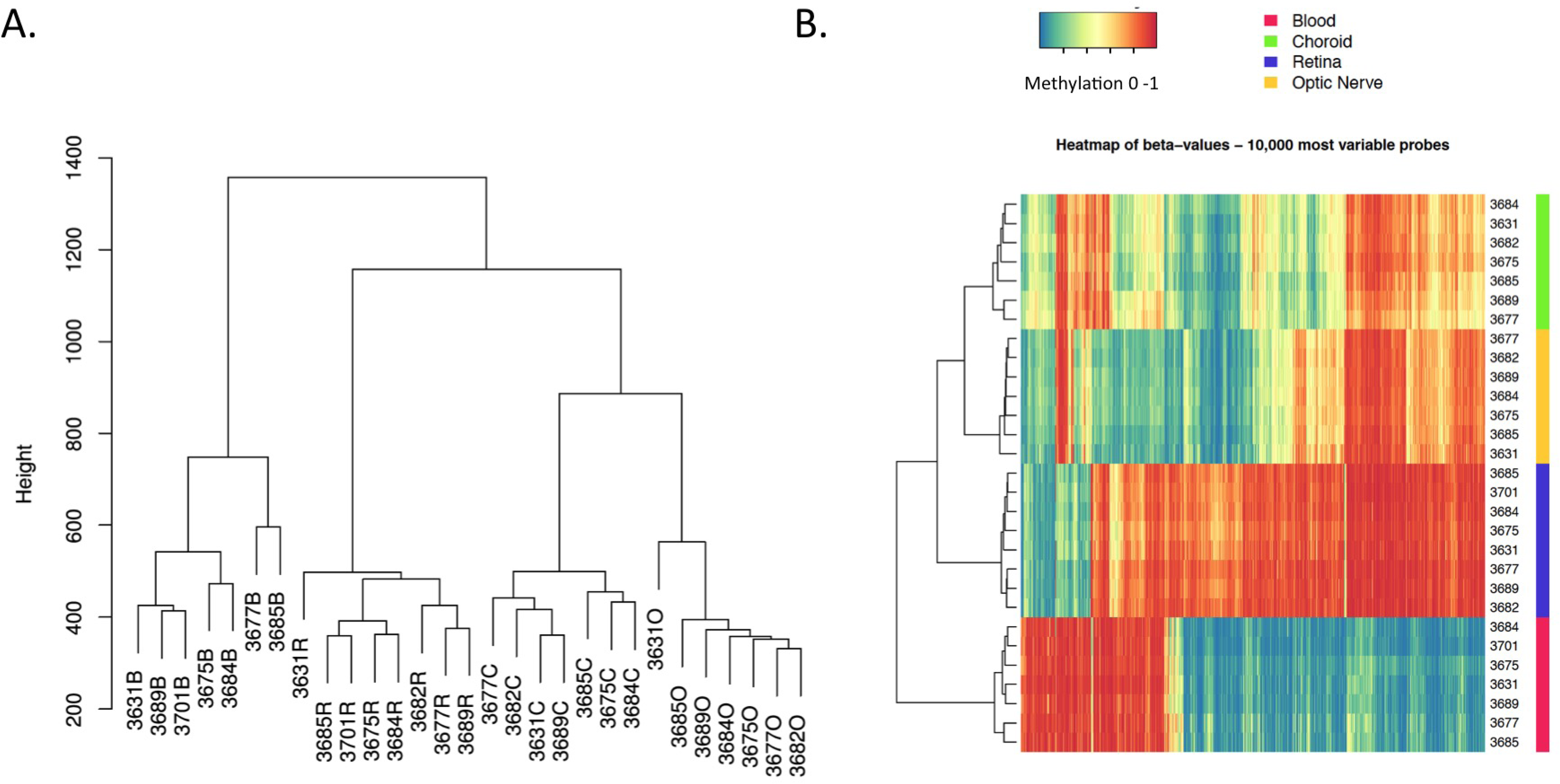
Sample relationships based on methylation at 433,768 HM450K probes shows tissue of origin as the main driver of variability within the dataset. (A) Hierarchical clustergram across all samples. Individuals are represented by their corresponding code followed by B (blood), C (RPE/choroid), R (retina), or O (optic nerve) to identify tissue type. (B) Heatmap of the 10,000 most variable HM450K probes. Probes are plotted on the x-axis, and individual tissue samples are plotted on the y-axis. Completely unmethylated (0 or 0% methylation) probes are represented as blue, and completely methylated (1 or 100% methylated) probes are represented as red.

To further dissect the source(s) of methylation variation within the dataset, we performed principal component analysis. As expected, most of the variation within the dataset (55.84%) was due to tissue type, with the first four principal components (PCs) all associated with tissue of origin at p < 0.05 (Figure 2). The only PC of variation that exclusively associated with individuals at p < 0.05 was PC13, comprising 802 probes that account for 1.87% of the variance within the dataset (**Supplementary Table 2**). Hierarchical clustering using probes associated with PC13 showed a clear separation of most individuals as predicted, despite very similar methylation levels across individuals overall (**Supplementary Figure 2**). Correlation analysis confirmed that PC13 was not related to any measured phenotype, such as age at death and cause of death (not shown).

**Figure 2.**
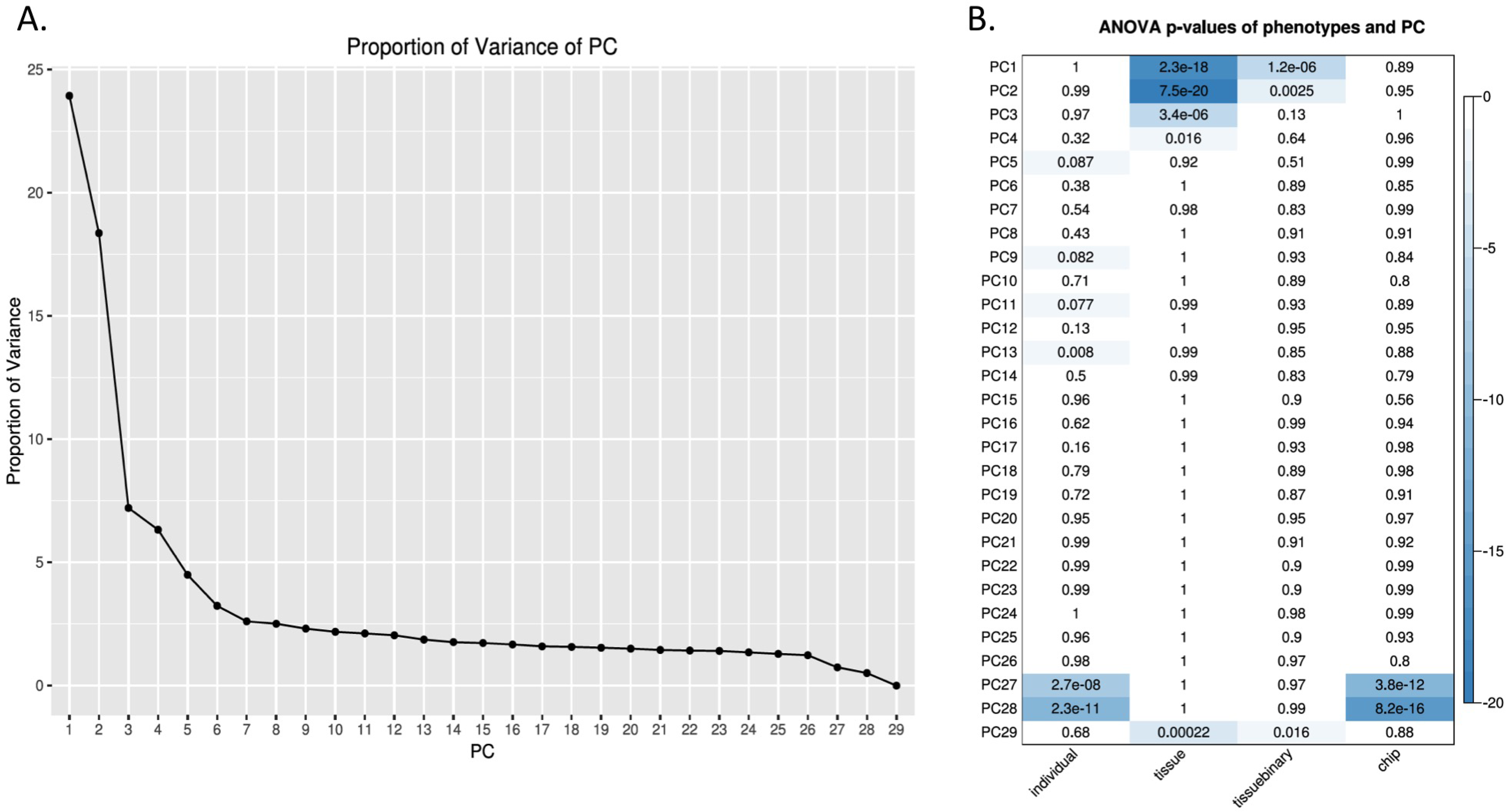
ANOVA analysis of principal components (PC) with sample traits. (A) Scree plot showing the proportion of total variation in the dataset captured by each of the first 29 PC. (B) Significance of association between each PC and individual, tissue of origin, eye vs non-eye (tissuebinary), and array chip. The first 4 PCs show a significant association with tissue of origin as anticipated, whereas PC13 captures the majority of inter-individual variation.

### Correlation of methylation values within and between tissues

Despite this discrete clustering, and reflecting the large number of CpG sites analysed, there was generally a strong correlation between methylation across all samples, with a median Spearman correlation of 0.914 (range: 0.819 – 0.987). Blood methylation profile had the strongest correlation with RPE/choroid (median r = 0.913, range: 0.892 – 0.939; not shown), followed by optic nerve (median, range) (not shown) and retinal tissue (median r = 0.850, range: 0.819 – 0871; not shown),.

Methylation β-values were bimodally distributed, with the majority of sites being commonly hypomethylated (β ≤ 0.2) or hypermethylated (β ≥ 0.8) across all tissues (**Supplementary Figure 3**). In order to explore this further, we divided probes into three categories of hypo (β ≤ 0.2), intermediate (0.2 < β <0.8) and hypermethylated (β ≥ 0.8) and then assessed the degree of overlap in each class of probe across all tissues tested. This approach confirmed that the majority of methylation is relatively consistent across all tissues, though considerable evidence of tissue-of-origin methylation levels is also present within all 3 broad methylation classes (Figure 3). Interestingly, each of the ocular tissues examined also shared a uniquely overlapping set of methylation levels with blood rather than other eye tissues (Figure 3).

**Figure 3.**
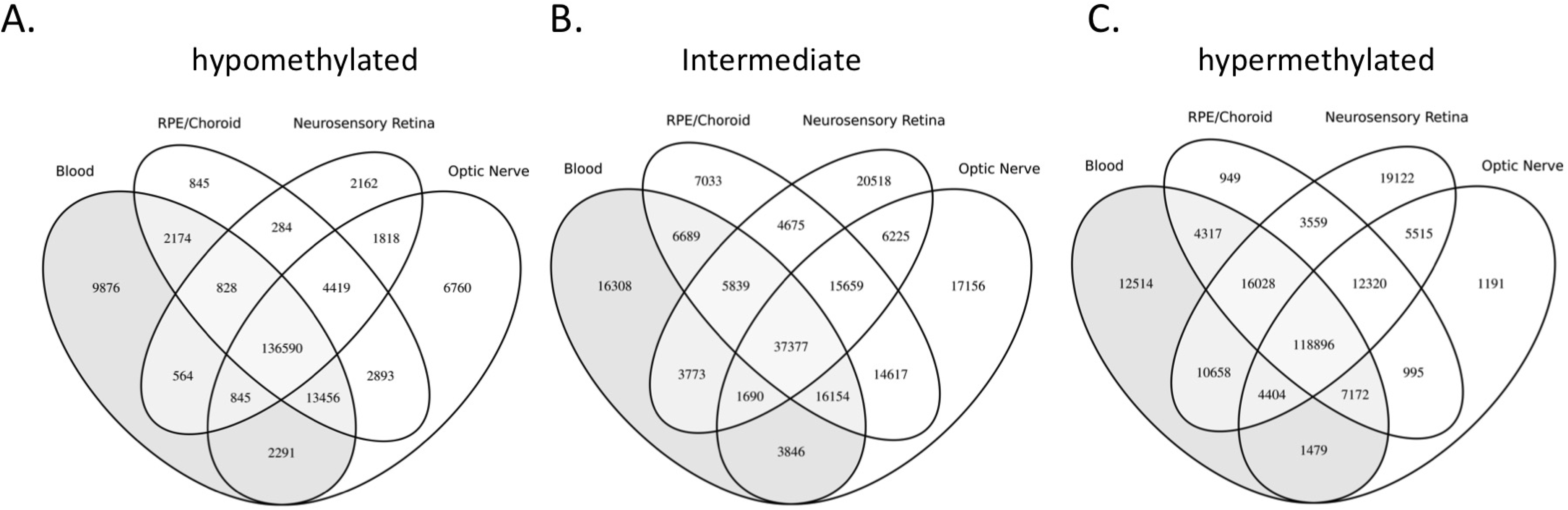
Venn diagrams showing the overlap between tissues of probes designated hypomethylated (<20%; A), intermediate (20–80%; B) and hypermethylated (>80%; C). Whereas the majority of probes showing conserved methylation across tissues are either hyper– or hypomethyalated, tissue specific methylation is more likely to be at intermediate levels.

We were primarily interested in identifying CpG sites where inter-individual variation in blood was also associated with inter-individual differences in eye methylation. Such probes may represent reliable proxy measures for variation in ocular methylation, measurable in peripheral blood. To investigate this, pairwise correlation analysis was performed for each probe within a subset identified as variable in blood across individuals, defined as those for which the inter-individual methylation difference between the 10^th^ and 90^th^ percentiles was > 5% (224,417 probes in total). The distribution of correlations between matched samples was compared with a randomly permuted distribution (i.e. null) to establish the degree of relatedness between blood and eye methylation levels. Significant differences were found between the true and null distributions (Wilcoxon rank test p < 2.2 x10^-16^ for all tissues (not shown)). Interestingly, most of the probes that were correlated or anti-correlated (Spearman r > 0.5, p < 0.05) between blood and eye tissues within an individual were unique for each pairwise tissue comparison, with only 122 probes commonly correlated with blood across all three eye tissues. Hierarchical clustering using these commonly variable probes revealed that retina and blood samples still cluster primarily by tissue, whereas RPE/choroid and optic nerve samples cluster by individual as opposed to tissue type (**Supplementary Figure 4**).

Thus, these probes in isolation are of limited utility in inferring general ocular methylation status from blood.

### Genomic context of variable probes

Compared to the overall distribution of probes, those showing inter-individual variation in blood were enriched for intergenic regions and CpG ‘open seas’ (Figure 4). The genomic distribution of PC13-related probes appeared to be closer to the overall distribution, rather than blood-variable probes, being enriched at sites within 200bp of transcription start sites, and also in CpG islands. This suggests that probes that vary between individuals do not necessarily co-vary between tissues types within an individual. Of 802 probes associated with PC13, 281 showed variable methylation in blood of different individuals, and of those, 115 showed correlated methylation with one or more eye tissues within an individual.

**Figure 4.**
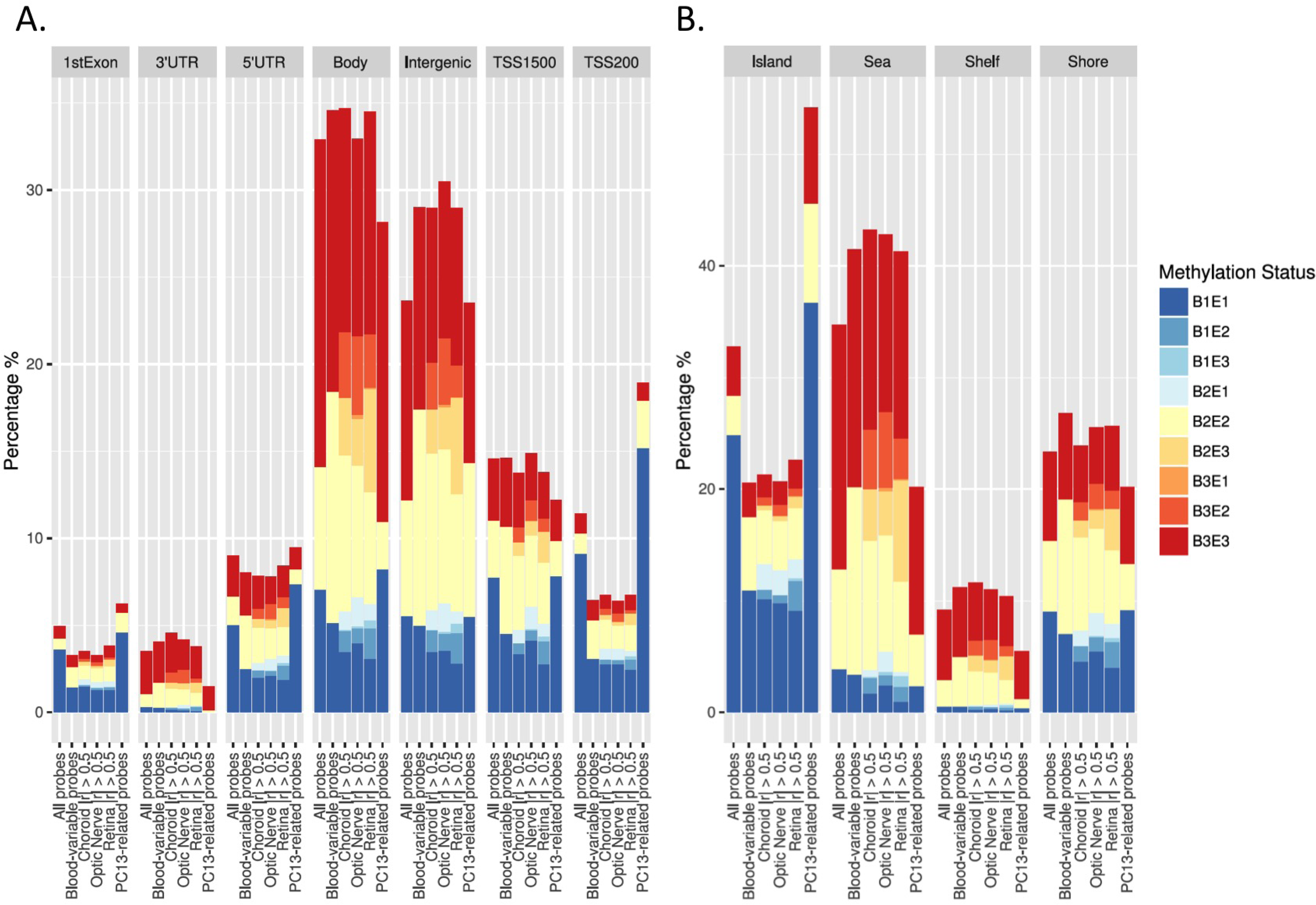
Distribution of mean methylation according to (A) genomic feature or, (B) CpG density relative to CpG islands. Classes B1, B2 and B3 correspond to hypomethylated, intermediate and hypermethylated probes in blood with E1, E2 and E3, the equivalent categories in eye tissue. (A) The majority of probes are located in gene bodies or intergenic regions and show hyper– or intermediate methylation in both eye and blood. In contrast, a larger proportion of probes in the 5’UTR, 1^st^ Exon or TSS regions are hypomethylated in both blood and eye tissues. {B) As anticipated, the majority of CpG island-associated probes are hypomethylated in blood and eye tissue, whereas CpG island Shores, Shelves and Open Sea are more highly methylated in both tissues. Data are shown for (i) all probes, (ii) probes showing inter-individual variation in blood, (iii – v) blood variable probes highly correlated with RPE/choroid, optic nerve or retina, and (vi) PC13-specific probes (all on the X-axis). The latter are primarily located within 200bp of gene transcription start sites in CpG island regions.

### Gene Pathway and ontology Analysis

The 802 probes associated with PC13 were linked to 541 genes. IPA analysis identified an overrepresentation of canonical pathways, ‘IL-1 Signaling’, ‘Molecular Mechanisms in Cancer’, ‘Protein Kinase A, Relaxin and α–Adrenergic signaling’. Gene Ontology analysis identified enrichment of cell cycle regulators, protein modifiers (particularly kinases) and genes involved in RNA metabolism. There was no enrichment for specific pathways in the genes associated with the 115 probe subset that showed a correlation with one or more eye tissue.

## DISCUSSION

Much of the application of epigenome-wide association studies (EWAS) to human conditions in recent years has been predicated on the assumption that accessible tissues show evidence of epigenetic variation associated with an exposure of interest, or reflective of less accessible tissues directly implicated in specific phenotypes. Exceptions to this include the use of tissue biopsies from adults (eg. adipose or muscle in metabolic conditions), tissue removed during surgery for clinical purposes (heart surgery and cardiovascular disease) and the use of tissues obtained postmortem (particularly of CNS origin). The latter are particularly important in allowing the delineation of the baseline epigenetic profile of tissues not accessible in living individuals and for comparing such a profile to more accessible tissues such as blood or saliva.

The potential for epigenetic variation to play a role in diseases of the eye has been suggested for several years, with various explorations in blood or saliva in individuals with conditions such as macular degeneration or retinitis pigmentosa. Recently, the potential for DNA methylation to play a role in Age-related Macular Degeneration (AMD) has been directly investigated through the genome-wide DNA methylation profiling of blood from AMD patients and controls. Small but replicable differences in DNA methylation were identified in the blood of neovascular AMD patients near the age-related maculopathy susceptibility 2 (*ARMS2*), gene, previously linked to AMD in genome-wide association studies (GWAS). Interestingly, an integrated analysis of blood and retina further identified consistent DNA methylation variation in the protease serine 50 (*PRSS50*) gene [14].

Genetic variation at methylation quantitative trait loci (mQTLs) plays a major role in shaping the human DNA methylation profile, including the potential modulation of the effects of environmental exposures on methylation variation. Although some mQTLs are likely tissue specific in their influence on methylation, others are likely conserved across tissues with their effects measureable in tissues such as blood. Evidence in support of a combined effect of genetic and epigenetic variation has also been implicated in AMD with evidence of an association with underlying genotype of the risk SNP rs10490924 with disease-associated variation in *ARMS2* methylation [14].

In many large epidemiological studies, whole blood is the only biological material that has been archived. The extent to which the DNA methylation patterns of easily accessible tissues like whole blood represent the epigenetic phenotype in inaccessible tissues is unclear, although recent studies are showing a certain degree of concordance. Epigenetic variation arising before disease could be inherited and thus be present in all adult tissues, or it could arise stochastically during the lifecourse and be limited to one, or a few tissues. Epigenetic variation can also be environmentally induced by life-style related factors, such as diet or smoking. In support of this, a panel of metastable epialleles has recently been described in humans that show inter-individual variation apparently independent of tissue of origin [15]. Such loci have considerable potential to act as epigenetic ‘sensors’ of past environmental influence/exposure.

In the current study, we defined the DNA methylation profile of 3 ocular tissues and compared these directly to matched peripheral whole blood from the same individuals. Despite a largely concordant methylation profile, only limited instances of correlated variation between tissues within specific individuals were observed. Collectively, our data support recent findings in similar analyses of the brain that suggested EWAS approaches using whole blood for disorders of the brain are likely to be of only limited utility for discovery type approaches aimed at unravelleing disease aetiolgoy [8]. Nevertheless, as with our findings in the eye, there are a proportion of sites where inter-individual variation is correlated between whole blood and brain, highlighting the utility (albeit limited with the current platform) of using a blood-based EWAS to identify potential biomarkers of disease of more central tissues such as the brain and the eye.

There are some caveats to the use of peripheral blood leukocytes to study DNA methylation profiles. Firstly, like neurosensory retinal and RPE/choroidal tissue, peripheral blood comprises a heterogeneous cell population. Therefore, any observed variation in DNA methylation assessed in whole blood may reflect small changes in the proportions of differentially methylated cell types that may in turn vary between individuals. A further limitation involves the issue of timing. It is now apparent that the DNA methylation of whole blood is altered with ageing and therefore samples taken from the same individual might be different if taken a few years apart. Equivalent data are generally not available for more central tissues of CNS or ocular origin. This has considerable potential to confound interpretation of EWAS studies.

## List of abbreviations

CV: coefficient of variation
EWAS: Epigenome-wide association study
DMP: differentially methylated probes
GO: Gene Ontology
GWAS: Genome-wide association study
HM450K: Illumina Infinium HumanMethylation450 BeadChips
RPE: retinal pigmented epithelium
SNP: single nucleotide polymorphisms

## Authors’ contributions

AWH, JJW, JEC and RS designed this study. Sample collection and processing were performed by AWH and MF, whilst JEJ performed the bisulfite conversion. Data were analysed by AWH, VJ, JEJ, HL and RS. This manuscript was prepared by AWH, VJ, JEJ, ASO and RS and all authors reviewed and gave final approval.

## Acknowledgments and Financial Support

We are grateful to Mrs Lisa Buckland and the staff of the Lions Eye Bank who facilitated sample collection. This work was supported by an Australian National Health and Medical Research Council (NHMRC) grant [APP1024619], the Ophthalmic Research Institute of Australia, the American Health Assistance Foundation and the Victorian Government’s Operational Infrastructure Support Program. RS is supported by a NHMRC Senior Research Fellowship and AWH by a NHMRC Early Career Fellowship.

## Data respository

All raw and processed files can be accessed through the GEO:/ArrayExpress Accession number (######).

## Figure and Table Legends

**Supplementary Figure 1.** Multi Dimensional Scaling plot showing the relationship between samples based on 1000 most variable probes. Whereas RPE/choroid (C) and optic nerve (O) tissue cluster together, with methylation profiles similar to adult cortex tissue, retinal (R) methylation profile is distinct, with blood (B) being least similar to all other tissues. A neuronal cell line is included for comparative purposes.

**Supplementary Figure 2.** (A) Heatmap of probes associated with PC13 at |r| > 0.5, showing some clustering according to individual rather than tissue type. Blue denotes fully unmethylated (0 or 0% methylation), while red fully methylated (1 or 100% methylated) probes. (B) Venn diagram showing the overlap of PC13-associated probes with each of the 3 ocular tissues.

**Supplementary Figure 3.** Distribution of mean HM450K methylation ⌷ –values (in 5% bins) across each tissue tested in the current study. All tisseus show a bimodal distribution. X-axis ⌷ – values correspond to methylation levels approximating 0–100%. Y-axis is the frequency of probes within each bin.

**Supplementary Figure 4.** Heatmap of 122 probes showing inter-individual variation in blood that show a degree of correlation between blood and all three eye tissues within individuals. Whereas clear inter-individual differences are apparent in the choroid and optic nerve, blood and retina remain clustered by tissue type. Blue denotes fully unmethylated (0 or 0% methylation), while red fully methylated (1 or 100% methylated) probes.

